# Nicotinic acetylcholine receptor signaling regulates cytokine production in Jurkat T cells

**DOI:** 10.1101/2025.09.24.678140

**Authors:** Bailey Laforest, Alain Simard

## Abstract

Nicotinic acetylcholine receptors (nAChRs) regulate immune cell functions, yet their expression patterns and roles in human T cells remain incompletely defined. The immunomodulatory effects of α7 nAChR signaling in human immune cells is complicated by the presence of the human-specific duplicated α7 (dupα7) subunit. Here, we investigated the expression and function of nAChRs in human Jurkat T cells, focusing on α7, dupα7, α9, and α10. Quantitative PCR revealed transcripts for all four subunits. Mitogenic stimulation with PMA, ionomycin, and ConA significantly downregulated α7, α9, and α10 expression while upregulating dupα7, suggesting dynamic remodeling of receptor composition during T cell activation. Functional assays showed that α7 antagonism with ArIB[V11L,V16D] strongly suppressed mitogen-induced IL-2 and TNF-α secretion, while nicotine pretreatment produced more modest reductions. Flow cytometry confirmed a decreased frequency of IL-2+ cells following treatment with nicotine or nAChR antagonists. These findings establish Jurkat cells as a tractable model for studying nAChR signaling in human T cells. Our results demonstrate that α7-containing nAChRs positively regulate cytokine production, while dupα7 expression increases during activation and may act as a negative regulator of α7 function. Together, these data highlight nAChRs as key modulators of T cell activity and identify α7 and dupα7 as potential therapeutic targets for regulating adaptive immunity.

## 1 Introduction

The nervous and immune systems are intricately connected through bidirectional communication, mediated by shared signaling molecules such as neurotransmitters and cytokines^1^. Among these, acetylcholine (ACh) is traditionally recognized as a neurotransmitter but is also produced and released by immune cells, creating a non-neuronal cholinergic network that modulates immune function^2,3^. CD4+ T cells, in particular, have robust choline acetyltransferase expression and ACh content, and their cholinergic activity is dynamically induced by T cell receptor (TCR) engagement and mitogenic stimulation^4–10^. They also express multiple nicotinic acetylcholine receptor (nAChR) subtypes, allowing them to directly respond to cholinergic cues^11–16^.

nAChRs are pentameric ligand-gated ion channels that bind to ACh and exogenous nicotine. They are composed of various combinations of α (α2-α7, α9, α10) and β (β2-β4) subunits, forming diverse heteromeric and homomeric receptor subtypes^17^. While best known for their neuronal functions, many nAChR subunits–including α7, α9, and α10–are expressed in immune cells^11–16^. The α7 subunit forms homomeric receptors, whereas α9 and α10 typically assemble into heteromeric complexes (**Figure 1**). Notably, α9α10 receptors are uniquely inhibited by nicotine, distinguishing them from other nAChR subtypes^18^. Activation of α7 nAChRs on macrophages has been shown to suppress inflammation via the cholinergic anti-inflammatory pathway^19,20^, while genetic deletion or selective antagonism of α9α10 nAChRs has also been linked to anti-inflammatory effects.

**Figure 1.**
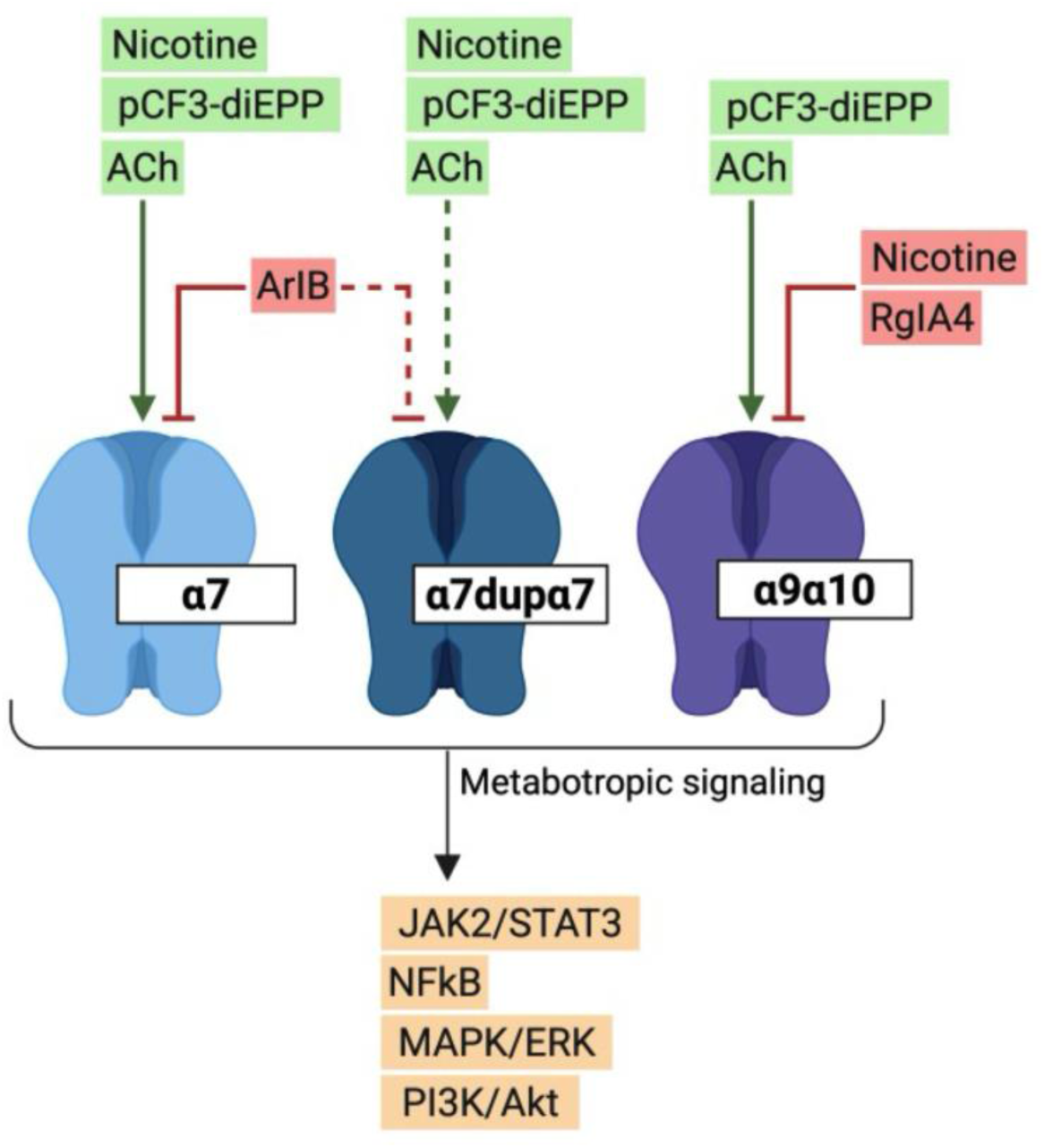
Ligands and metabotropic signaling of nAChR subtypes. The α7, dupα7, and α9α10 nAChR subtypes respond to acetylcholine (ACh) and selective exogenous ligands. Nicotine activates α7 receptors but inhibits α9α10 receptors^18^, while the silent agonist pCF3-diEPP can engage α7/α7dupα7, and α9α10^42^. The α7 antagonist ArIB blocks α7/α7dupα7-mediated responses, whereas the α9α10 antagonist RgIA4 blocks α9α10-mediated responses^43–45^. Downstream, nAChRs couple to intracellular signaling cascades including JAK2/STAT3, NFκB, MAPK/ERK, and PI3K/Akt, highlighting the non-canonical, metabotropic roles of nAChRs in cholinergic immunoregulation^35,36^. Image created with Biorender.

In T cells, cholinergic signaling influences intracellular Ca^2+^ flux, proliferation, cytokine production, and differentiation^21–29^. In rodent models of autoimmunity and chronic inflammation, nicotine treatment consistently improves disease outcomes, primarily by modulating autoreactive T cell activity and proinflammatory cytokine production^30–33^. Although these findings highlight nAChRs as key regulators of T cell responses, the specific roles of individual receptor subtypes remain unresolved.

Unlike their classical ionotropic functions in neurons, nAChRs in immune cells primarily signal through non-canonical metabotropic mechanisms^34^. These involve direct interactions of intracellular receptor domains with signaling proteins, activating cascades such as JAK2/STAT3, PI3K/Akt, and MAPK/ERK (**Figure 1**)^35,36^. For example, in T cells, α7 can form complexes with the TCR/CD3 complex and couple to protein tyrosine kinases, mobilizing intracellular Ca^2+^ independently of ion channel activity^23^. Such mechanisms challenge assumptions derived from neuronal electrophysiology and may explain the unconventional effects of many nAChR ligands in immune cells^37,38^.

Various pharmacological tools, including silent agonists and conopeptides, have been instrumental in dissecting nAChR signaling (**Figure 1**)^39,40^. Silent agonists bind to nAChRs without inducing classical ion currents but stabilize conformations that support metabotropic signaling^41^. This property allows them to modulate intracellular pathways selectively, making them useful probes for non-conducting receptor functions. Recently, the α7- and α9α10-selective silent agonist, pCF3-diEPP, was shown to reduce the production of proinflammatory cytokines from human immune cells^42^.

Conopeptides, such as ArIB[V11L,V16D] (ArIB) and RgIA4, are highly specific antagonists of α7 and α9α10 nAChRs, respectively^43–45^. Their subtype specificity allows precise interrogation of receptor contributions to cellular responses, and they have been widely used to distinguish the overlapping functions of nAChR subtypes in neuronal and immune contexts^42,46^. Together, these ligands provide powerful tools for exploring the subtype-specific role of nAChRs in T cells.

A unique aspect of cholinergic signaling in human biology is the presence of *CHRFAM7A*, a human-specific partial duplication of the α7 gene (*CHRNA7*), which encodes the dupα7 subunit^47–49^. Unlike α7, dupα7 cannot form functional homomeric receptors but assembles with α7 to generate dominant-negative receptors that dampen α7-mediated signaling^50–53^. Dupα7 is highly expressed in immune cells, suggesting a specialized role in immune modulation^54–59^. Failure to account for dupα7 in experimental design and interpretation likely contributes to translational gaps between rodent models (which lack dupα7) and human studies^60^. Thus, distinguishing ɑ7 from dupɑ7 at the transcript and protein levels is critical for understanding cholinergic regulation of human immunity.

Within this evolving landscape, several key knowledge gaps persist. First, the baseline and activation-induced expression of nAChR subunits in human T cells has not been consistently quantified with resolution of ɑ7 versus dupɑ7. Second, whether nAChR signaling enhances or restrains T cell cytokine production likely depends on receptor composition, ligand state, and ionotropic versus metabotropic signaling. Third, the effects of selective α7 and α9α10 ligands on T cell cytokine production have not been systematically compared.

To address these questions, we characterized the gene expression of α7, dupα7, α9, and α10 in Jurkat T cells at baseline and after mitogenic stimulation, and assessed how selective activation or inhibition of these receptors influences cytokine production. Our findings reveal that the T cell nAChR repertoire is remodeled during mitogen-induced activation and that nAChR signaling supports cytokine production. These results refine mechanistic models of cholinergic immunoregulation, highlight the significance of dupα7 in human T cell biology, and suggest receptor-subtype-specific strategies for modulating T cell-mediated inflammation.

## 2 Materials and Methods

### 2.1 T Cell Culture and Treatment

Jurkat E6-1 T cells (ATCC TIB-152) and CCRF-CEM T cells (ATCC, CCL-119) were maintained in RPMI-1640 (ATCC, #30-2001) supplemented with 10% fetal bovine serum (Gibco, #12484028) and 1% penicillin-streptomycin (Gibco, #15140122) at 37°C with 5% CO2 (Forma Series II Water Jacketed CO2 Incubator, ThermoFisher).

For Jurkat qPCR experiments, cells were seeded at 5×10^5^ cells/mL in T25 flasks (5 mL/flask) and stimulated for 2 h or 48 h with various combinations of phorbol 12-myristate 13-acetate (PMA; 50ng/mL; Sigma, #P8139), ionomycin (Iono; 1.5 μM; Tocris, #1704), and Concanavalin A (ConA; 5 μg/mL; MP Biomedicals, #195283), collectively referred to as PIC. For CCRF-CEM qPCR experiments, cells were seeded at 8×10^5^ cells/mL in T25 flasks (5 mL/flask) and treated for 6 h, 24 h, or 48 h with various combinations of ConA (3 μg/mL), nicotine (10 μM), pCF3-diEPP (100 μM), ConA + nicotine, or ConA + pCF3-diEPP.

For cytokine assays, Jurkat cells were seeded at 5×10^5^ cells/mL in 24-well (1 mL/well) or 12-well (2 mL/well) plates for ELISAs and flow cytometry, respectively, and pretreated for 1 h with combinations of the following nAChR ligands: nicotine (100 μM; Tocris, #3546), pCF3-diEPP (100 μM; α7/α9α10 silent agonist), ArIB (500 nM; α7 antagonist), and RgIA4 (200 nM; α9α10 antagonist). Cells were then stimulated with PIC for 5 h (TNF-α) or 24 h (IL-2). Controls received an equivalent volume of DMSO (0.1%). pCF3-diEPP (para-trifluoromethyl-N,N-diethyl-N’-phenyl-piperazinium) was synthesized as previously described^61^. The conopeptides ArIB and RgIA4 were synthesized and characterized as previously described^45,62,63^.

### 2.2 Quantitative PCR

RNA was extracted using the PureLink RNA Mini Kit (Invitrogen, #12183025) with on-column DNase treatment (Invitrogen, #12185010), following the manufacturer’s instructions. RNA concentration and purity were assessed by spectrophotometry (NanoDrop, Thermo Fisher). cDNA was synthesized using the SensiFAST cDNA Synthesis Kit (Bioline, #BIO-65054) on an MJ Mini Thermal Cycler (Bio-Rad) with the following program: 25°C for 10 min, 42°C for 15 min, 85°C for 5 min. No-RT controls were prepared by replacing reverse transcriptase with water.

qPCR was performed with SYBR Green Master Mix (Applied Biosystems, #A25741) on a CFX Connect Real-Time PCR Detection System (Bio-Rad) using the following cycling conditions: 50°C for 2 min, 95°C for 2 min, then 40 cycles of 95°C for 3 s and 60–61°C for 30 s. No-template and no-RT controls were included for every target. Subunit-specific primers for α7 (*CHRNA7*), dupα7 (*CHRFAM7A*), α9 (*CHRNA9*), and α10 (*CHRNA10*) were designed and validated by Sanger sequencing. *HNRNPL* and *RER1* were used as reference genes due to their stable expression across diverse human tissues^64^. Relative expression was calculated using the 2^-ΔΔCt^ method. Primer sequences are listed in **Table 1**.

**Table 1.**
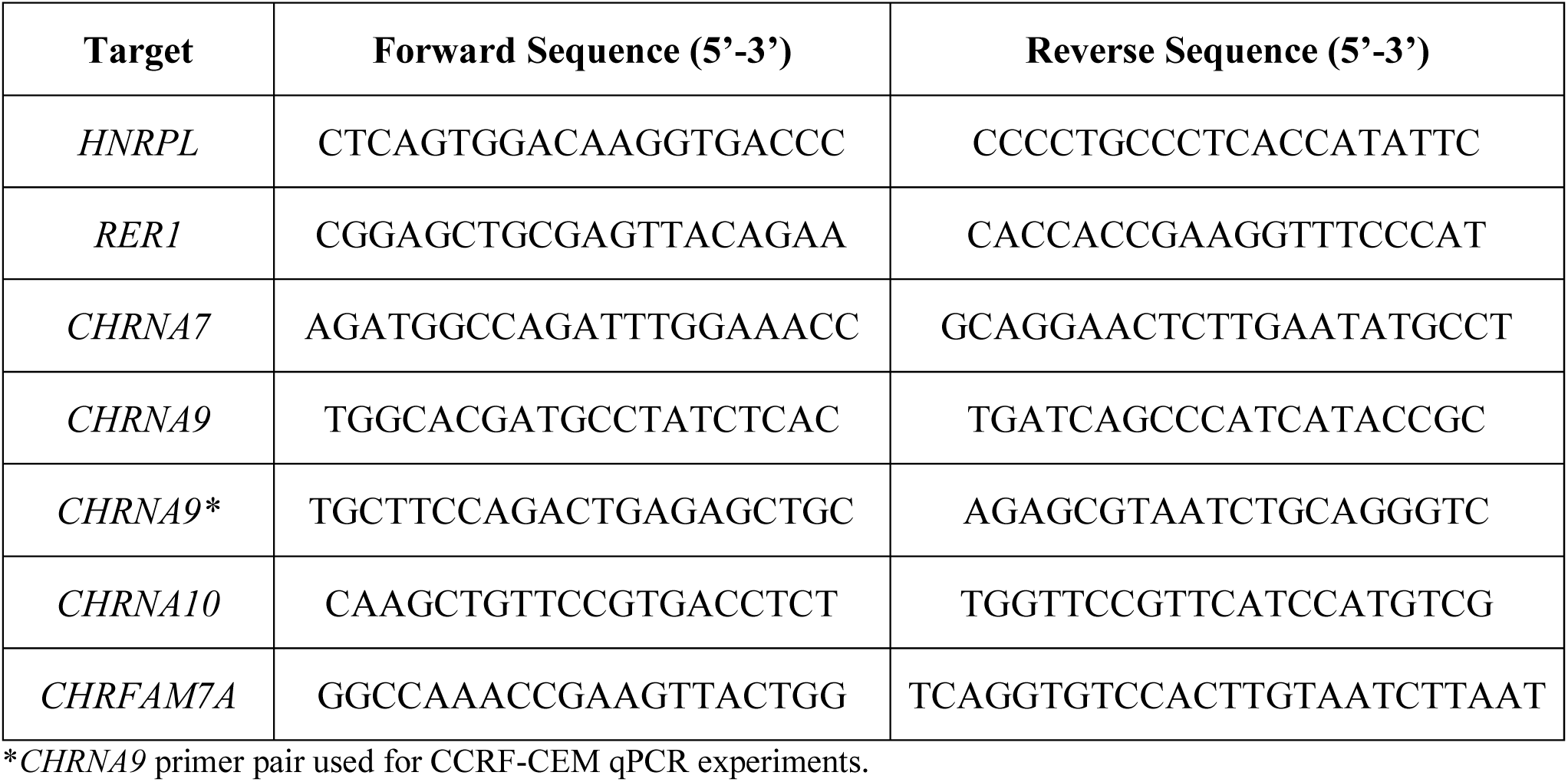
Primer sequences for qPCR.

### 2.3 PCR Product Purification and Sanger Sequencing

Endpoint Jurkat PCR products were purified with the PureLink PCR Purification Kit (Invitrogen, #K310002). DNA concentration and purity were assessed by NanoDrop One. Sequencing reactions contained ∼10 ng purified amplicon plus 50 ng forward or reverse primer in 7.7 µL. Sanger sequencing was performed at The Center for Applied Genomics DNA Sequencing Facility (SickKids). Chromatograms were inspected and aligned to NCBI RefSeq transcripts.

### 2.4 Cytokine Quantification by ELISA

Cell culture supernatants were collected at 5 h (TNF-α) and 24 h (IL-2) post-mitogenic stimulation. Cytokine concentrations were determined using human TNF-α (#DTA00D) and IL-2 (#D2050) Quantikine ELISA kits (R&D Systems) according to the manufacturer’s instructions. Samples were diluted 1:4 prior to IL-2 quantification. All samples were assayed in triplicate, and absorbance was measured at 450 nM with correction at 540 nM using a Synergy HTX Multimode Reader (BioTek).

### 2.5 Intracellular Cytokine Staining and Flow Cytometry

For intracellular cytokine detection, cells were treated as described above, and brefeldin A (5 μg/mL; BioLegend, #423101) was added during the final 5 h to block cytokine release. Cells (0.5-1× 10^6^/sample) were harvested, washed with PBS, and stained with Zombie Aqua viability dye (BioLegend, #423101) for 20 min at room temperature to exclude dead cells. After washing with flow buffer (1× PBS, 1% BSA, 2 mM EDTA), cells were stained with eFluor 450–anti-CD3 (Invitrogen, #48-0038-82) for 30 min at 4°C, then washed twice with flow buffer. Cells were fixed (Invitrogen, #00-8222-49) for 20 min at room temperature, washed twice with permeabilization buffer (Invitrogen, #00-8333-56), and stained with PE–anti–IL-2 (Invitrogen, #12-7029-81) and PerCP-Cy5.5–anti–TNF-α (Invitrogen, #45-7349-42) for 20 min at room temperature. After two washes with permeabilization buffer, cells were resuspended in 500 μL flow buffer.

Data were acquired on a SONY SA3800 Spectral Cell Analyzer (SA3800 Software v2.0.5.54250), with a minimum of 10,000 collected per sample. FlowJo software (v10.8.2) was used for analysis. Spectral references were generated from single-stained controls using Jurkat cells or compensation beads (Invitrogen, #01-3333-42). Heat-killed cells (10 min at 65°C) served as positive controls for Zombie Aqua staining. An unstained viable control was included to assess autofluorescence. Gating excluded debris (FSC vs SSC) and doublets (FSC-A vs FSC-H). Live cells were gated by Zombie Aqua negativity, and cytokine-positive gates were defined using untreated controls. Antibodies, fluorophores, clones, and working concentrations are listed in **Table 2**.

**Table 2.** Antibodies for flow cytometry.

| Target | Fluorophore | Clone | μL/100 μL |
| --- | --- | --- | --- |
| CD3 | eFluor 450 | UCHT1 | 2.5 |
| IL-2 | PE | MQ1-17H12 | 1.25 |
| TNF-α | PerCP-Cy5.5 | MAb11 | 5 |
| Viability | Zombie Aqua | - | 0.1 |

### 2.6 Statistical analysis

Data are presented as mean ± SEM. Statistical significance was determined by one-way or two-way ANOVA with Šídák’s post hoc test for multiple comparisons, as appropriate. A value of *p* < 0.05 was considered statistically significant. All analyses were performed using GraphPad Prism (v10.6.0). Biological replicates (n) are reported in figure legends and represents independent experiments performed with different cell passages.

## 3 Results

### 3.1 Mitogenic stimulation differentially regulates nAChR subunit expression in Jurkat cells

To determine whether nAChR subunit expression changes during T cell activation, Jurkat cells were stimulated with PMA, ionomycin (Iono), and ConA (PIC; **Figure 2A**). These mitogens provide a robust stimulus that mimics TCR activation^65^. PMA + Iono and ConA alone were also tested to isolate their individual effects. Untreated or DMSO controls served as baseline conditions.

**Figure 2.**
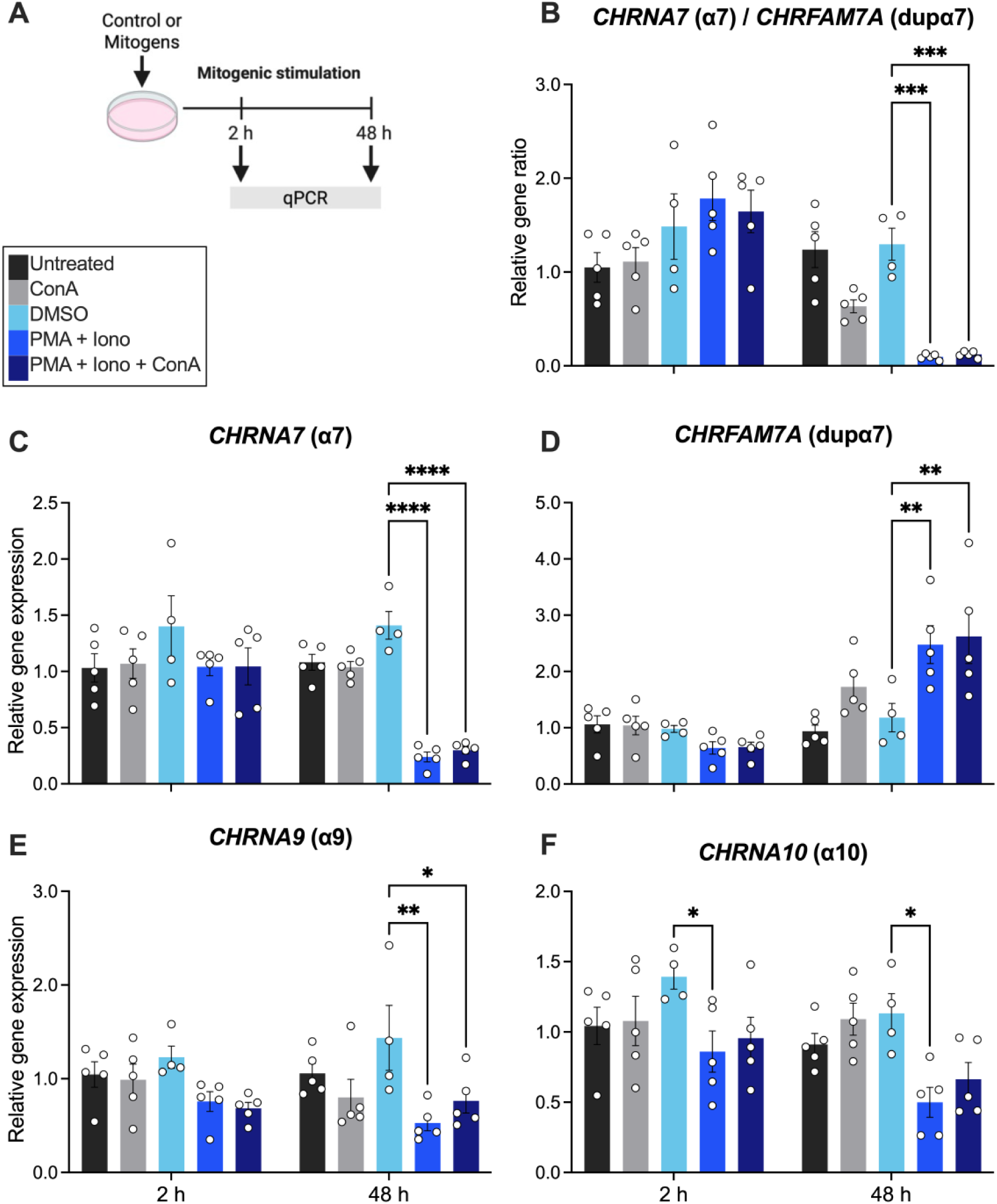
Mitogenic stimulation differentially regulates nAChR subunit expression in Jurkat T cells. Jurkat cells were treated for 2 or 48 h with the following conditions: untreated (black), ConA (5 μg/mL; gray), DMSO vehicle control (0.1%; light blue), PMA (50 ng/mL) + Iono (1.5 μM; medium blue), or PMA + Iono + ConA (dark blue). **(A)** Schematic of experimental design. Created with Biorender. **(B-F)** Gene expression was measured by qPCR and normalized to housekeeping genes. The 2^-ΔΔCt^ for each treatment is relative to 2 h untreated control. **(B)** *CHRNA7/CHRFAM7A* ratio, **(C)** *CHRNA7*, **(D)** *CHRFAM7A*, **(E)** *CHRNA9*, and **(F)** *CHRNA10*. Data are presented as mean ± SEM. Individual points represent biological replicates (n = 4-5/condition). Statistical significance was determined by two-way ANOVA with Šídák’s multiple comparisons test; *p < 0.05, **p < 0.01, ****p < 0.0001.

Jurkat cells expressed transcripts for α7, α9, α10, and dupα7 both at baseline and after mitogenic stimulation (**Figure 2C-F**). There was a significant interaction between treatment and time on α7 (*p* = 0.0004; **Figure 2C**) and dupα7 (*p* = 0.0001; **Figure 2D**), indicating that mitogenic stimulation altered these genes in a time-dependent manner. For α9, there was a significant treatment effect (*p* = 0.0008; **Figure 2E**), while α10 expression showed significant effects of both treatment (*p* = 0.0009) and time (*p* = 0.0166; **Figure 2F**). After 48 h of PMA + Iono stimulation, α7 (*p* < 0.0001), α9 (*p* = 0.0015), and α10 (*p* = 0.0119) were significantly downregulated, whereas dupα7 was upregulated (*p* = 0.0045), relative to DMSO controls. ConA alone had no effect on expression. α10 was also significantly reduced after 2 h of PMA + Iono treatment (*p* = 0.0477). Together, these data indicate that mitogenic stimulation shifts the α7/dupα7 balance toward higher dupα7 expression and lower α7 (**Figure 2B**), while also downregulating α9 and α10. This pattern may reflect a feedback mechanism that limits α7-dependent cholinergic signaling during sustained T cell activation.

For comparison, CCRF-CEM T cells were also assessed. These cells robustly expressed dupα7, α9, and α10, but showed very low α7 signal (Ct > 35) that was not reliably quantifiable by qPCR. There was a significant time effect on dupα7, α9, and α10 expression (p < 0.0001; **Figure 3A-D**). Over time, dupα7 and α9 increased, whereas α10 decreased, independent of mitogenic or cholinergic stimulation.

**Figure 3.**
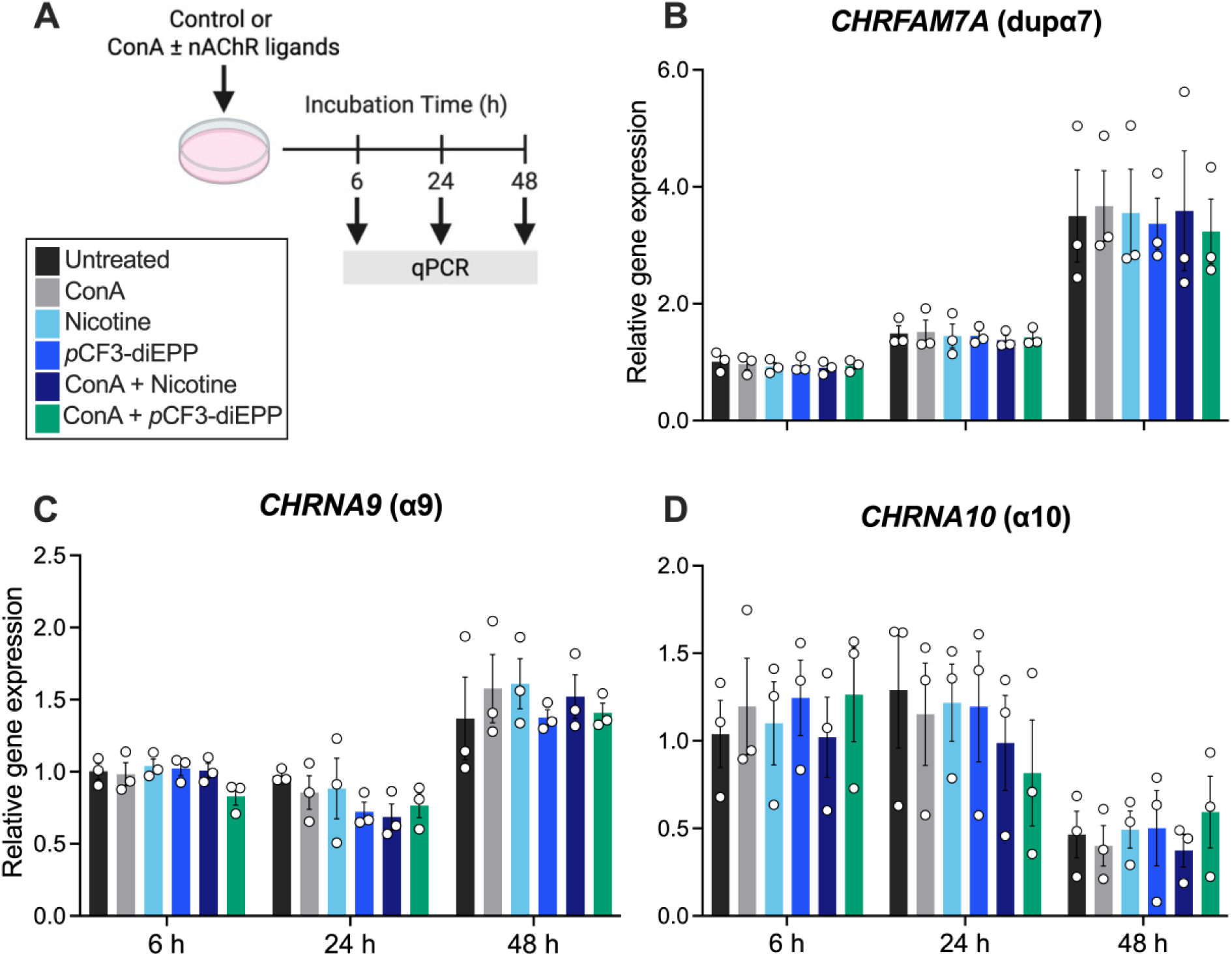
nAChR subunit expression is differentially regulated over time in CCRF-CEM T cells. CCRF-CEM cells were treated for 6 h, 24 h, or 48 h with the following conditions: untreated (black), ConA (3 µg/mL; gray), nicotine (10 µM; light blue), pCF3-diEPP (100 µM; medium blue), ConA + nicotine (dark blue), or ConA + pCF3-diEPP (green-blue). **(A)** Schematic of experimental design. Created with Biorender. **(B-D)** Gene expression was measured by qPCR and normalized to housekeeping genes. The 2^-ΔΔCt^ for each treatment is relative to 6 h untreated control. **(B)** *CHRFAM7A*, **(C)** *CHRNA9*, and **(D)** *CHRNA10*. Data are presented as mean ± SEM. Individual points represent biological replicates (n = 3/condition). Statistical significance was determined by two-way ANOVA with Šídák’s multiple comparisons test.

### 3.2 ɑ7 antagonism suppresses mitogen-induced cytokine production from Jurkat cells

To assess whether nAChR ligands modulate cytokine production, Jurkat cells were pretreated for 1 h with nicotine (100 μM) or the silent agonist pCF3-diEPP (100 μM), with or without the α7 antagonist ArIB (500 nM) or the α9α10 antagonist RgIA4 (200 nM), followed by PIC stimulation (**Figure 4A**). Unpretreated (PIC only) or DMSO-pretreated controls were also prepared. IL-2 was measured at 24 h and TNF-α at 5 h by ELISA. PIC stimulation robustly increased IL-2 (2.2–5.0 ng/mL) and TNF-α (24–55 pg/mL) secretion (**Figure 4B,C**). Nicotine alone reduced cytokine production but did not reach statistical significance. There was a significant overall effect of treatment, with ArIB alone significantly decreasing both IL-2 (*p* = 0.0309) and TNF-α (*p* = 0.0104). Nicotine combined with ArIB (*p* = 0.0314) or RgIA4 (*p* = 0.0454) also significantly reduced cytokine levels. In contrast, pCF3-diEPP, alone or with antagonists, had no effect. These findings suggest that α7 nAChRs positively regulate cytokine secretion in Jurkat T cells.

**Figure 4.**
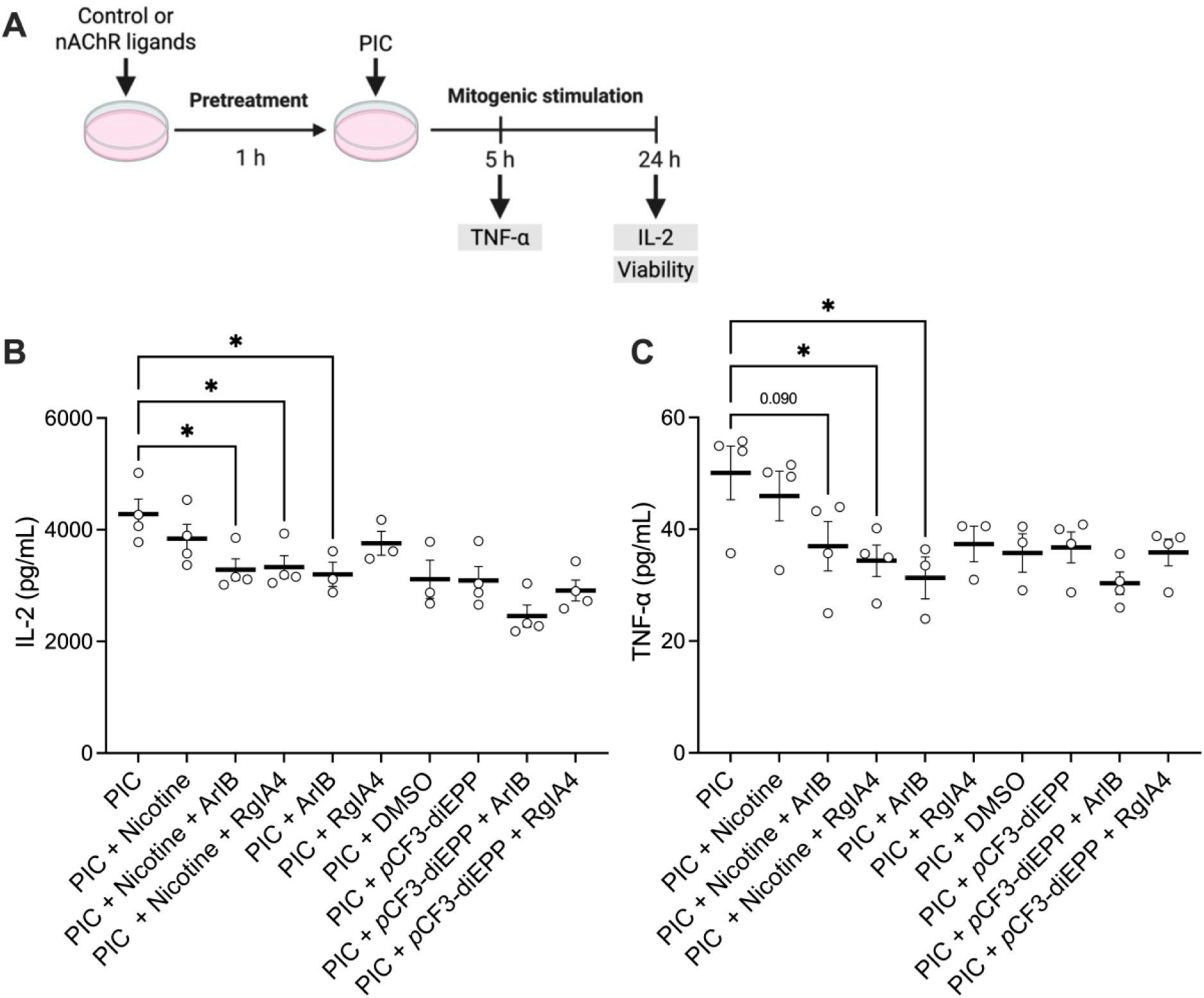
nAChR ligands regulate cytokine secretion from Jurkat T cells. Jurkat cells were pretreated for 1 h with various combinations of nicotine (100 µM), pCF3-diEPP (100 µM), the ⍺7-selective antagonist ArIB (500 nM), and the ⍺9/⍺9⍺10-selective antagonist RgIA4 (200 nM) prior to 24 h stimulation with PMA (50 ng/mL) + ionomycin (Iono; 1.5 µM) + ConA (5 µg/mL; PIC). Unpretreated (PIC only) and 0.1% DMSO-pretreated control groups were included. **(A)** Schematic of experimental design for all functional assays. Created with Biorender. The concentration of **(B)** IL-2 and **(C)** TNF-⍺ was quantified in cell culture supernatants using R&D Systems’ Quantikine ELISA kit. Data are presented as mean ± SEM. Individual points represent biological replicates (n = 3-4/condition). Statistical significance was determined by one-way ANOVA with Šídák’s multiple comparisons test; *p < 0.05.

### 3.3 nAChR signaling regulates the proportion of IL-2+ Jurkat cells

We next examined how nAChR ligands influence the percentage of cytokine-producing cells using flow cytometry (**Figure 5A-C**; see **Figure 4A** for experimental design). After 24 h of PIC stimulation, ∼11-13% of Jurkat cells were IL-2+; compared to <0.5% in untreated controls (**Figure 5A,B**). Nicotine pretreatment significantly reduced %IL-2+ cells (*p* = 0.0038), both alone and in combination with ArIB (*p* = 0.0135) or RgIA4 (*p* = 0.0096). pCF3-diEPP trended toward a reduction (*p* = 0.0958), but the effect became significant when combined with ArIB (*p* = 0.0255) or RgIA4 (*p* = 0.0281). CD3 expression decreased after stimulation, consistent with TCR internalization, but was unaffected by nAChR ligands (**Figure 5C**). TNF-α+ events were below the detection limit after 5 h and are not reported. Cell viability remained >95% across all treatments (**Figure 6A,B**), indicating that differences in cytokine production reflected immunomodulation rather than cytotoxicity.

**Figure 5.**
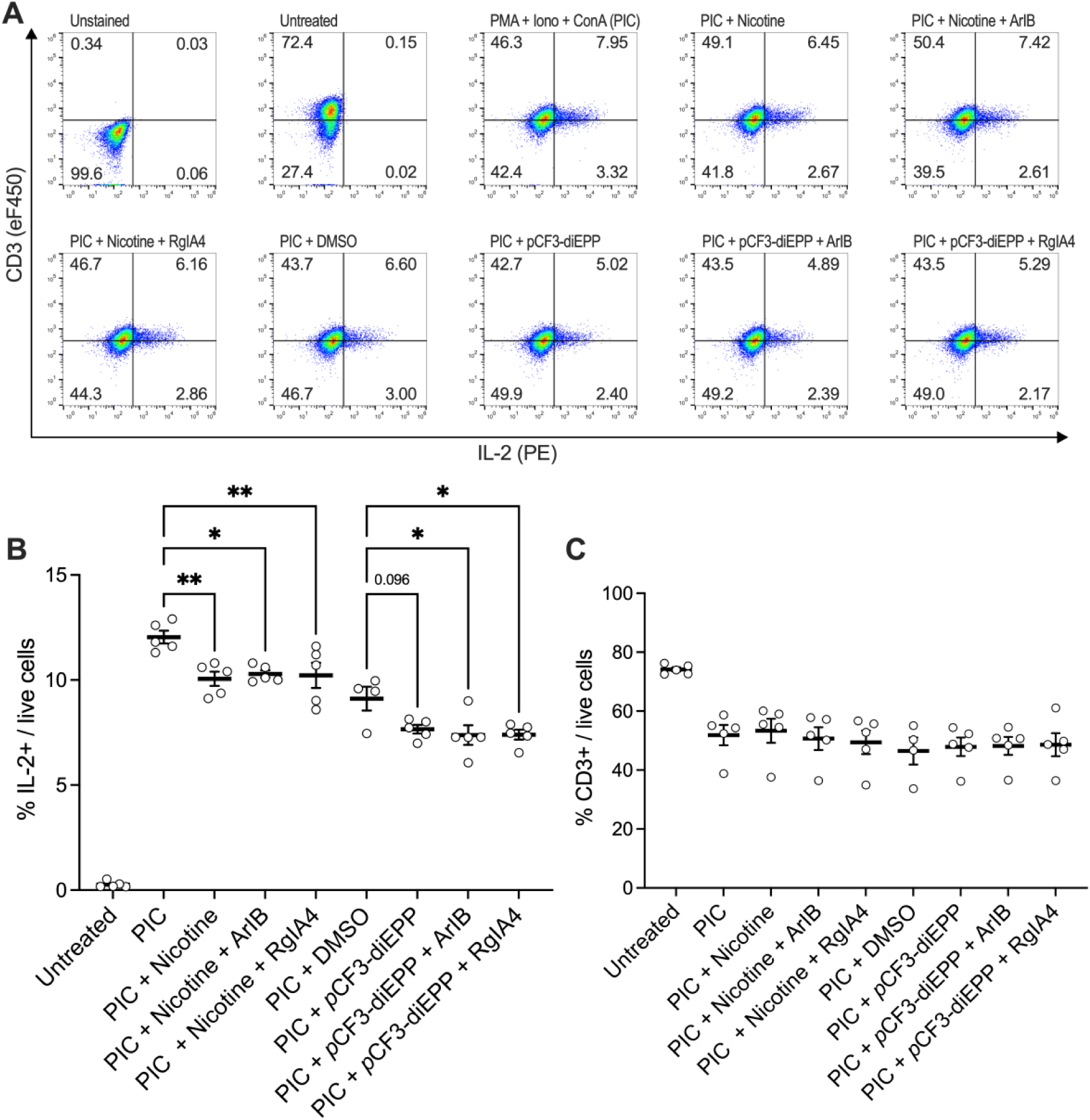
nAChR ligands regulate percentage of IL-2+ Jurkat T cells. Jurkat cells were pretreated for 1 h with various combinations of nicotine (100 µM), pCF3-diEPP (100 µM), the ⍺7 antagonist ArIB (500 nM), and the ⍺9⍺10 antagonist RgIA4 (200 nM) prior to 24 h stimulation with PMA (50 ng/mL) + Iono (1.5 µM) + ConA (5 µg/mL; PIC). See Figure 4A for experimental design. Unpretreated (PIC only) and DMSO (0.1%)-pretreated control groups were included (See Figure 4A for experimental design). Brefeldin A (BFA, 5 µg/mL) was added during the final 5 h to inhibit cytokine secretion. **(A)** Representative flow cytometry plots showing IL-2 vs CD3 staining for each condition. The percentage of **(B)** IL-2+ and **(C)** CD3+ cells were quantified by flow cytometry. Data are presented as mean ± SEM. Individual points represent biological replicates (n = 4-5/condition). Statistical significance was determined by one-way ANOVA with Šídák’s multiple comparisons test; *p < 0.05, **p < 0.01.

**Figure 6.**
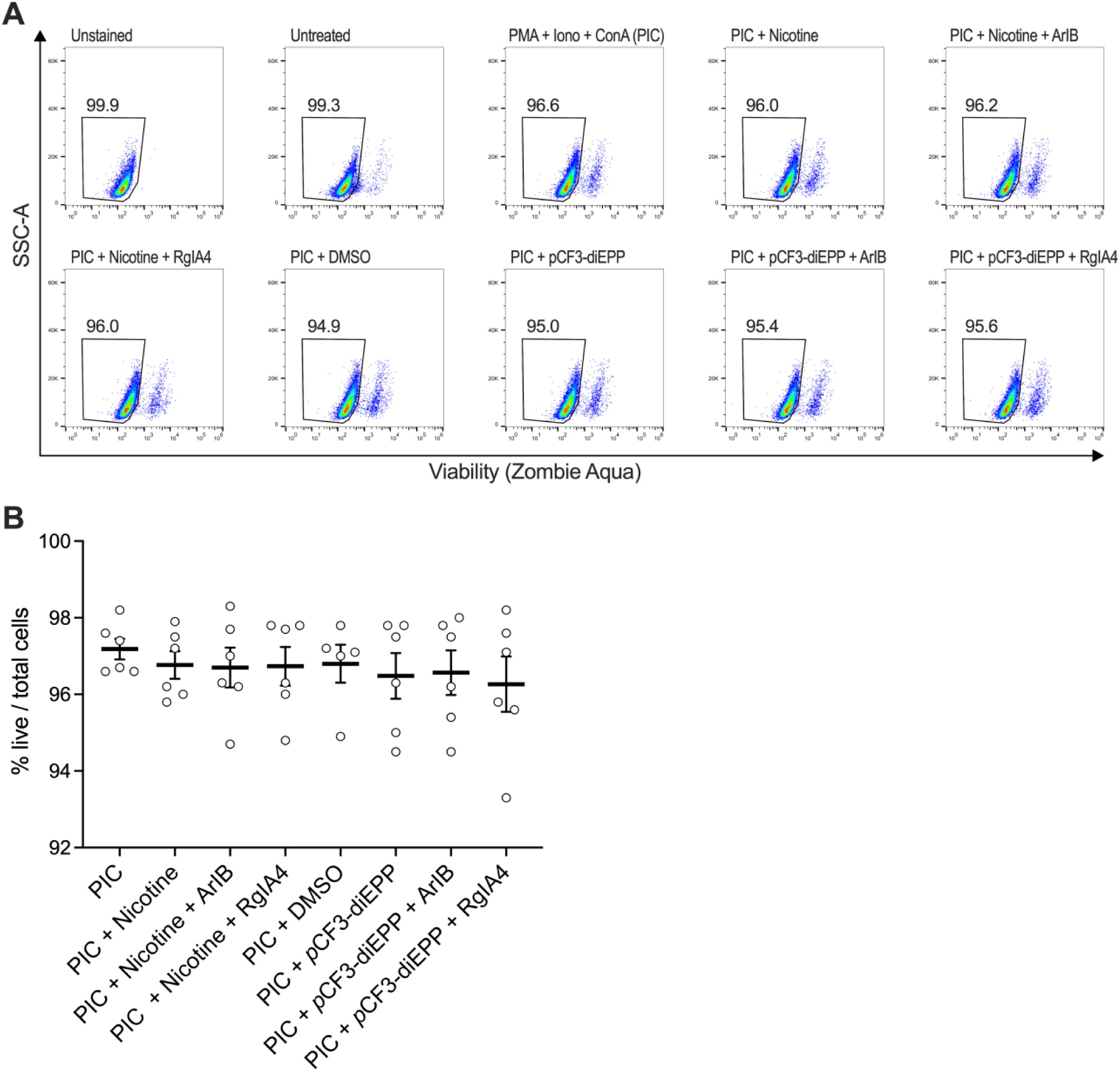
nAChR ligands and mitogenic stimulation do not affect Jurkat T cell viability. Jurkat cells were pretreated for 1 h with various combinations of nicotine (100 µM), pCF3-diEPP (100 µM), the ⍺7-selective antagonist ArIB (500 nM), and the ⍺9/⍺9⍺10-selective antagonist RgIA4 (200 nM) prior to 24 h stimulation with PMA (50 ng/mL) + ionomycin (Iono; 1.5 µM) + ConA (5µg/mL; PIC). Unpretreated (PIC only) and DMSO (0.1%)-pretreated control groups were included (See Figure 4A for experimental design). Brefeldin A (BFA, 5 µg/mL) was added during the final 5 h to inhibit cytokine secretion. **(A)** Representative flow cytometry density plots showing zombie aqua viability staining for each condition. The percentage of **(B)** zombie aqua negative cells were quantified by flow cytometry. Data are presented as mean ± SEM. Individual points represent biological replicates (n = 4-5/condition).

## 4 Discussion

This study provides new insights into the expression, regulation, and function of nAChRs in human T cells. Using Jurkat and CCRF-CEM T cells, we confirmed expression of α7, α9, α10, and the human-specific dupα7, and demonstrated that their expression is dynamically regulated by mitogenic stimulation. Pharmacological experiments revealed that nAChR ligands modulate cytokine production in complex, subtype-specific ways. Most notably, the α7 antagonist ArIB produced robust immunosuppressive effects, reducing both IL-2 and TNF-α secretion and lowering the percentage of IL-2+ cells.

Dupα7 mRNA was detected in both Jurkat and CCRF-CEM cells, consistent with previous reports that dupα7 is often the predominant transcript in human immune cells^48,53,54,59,66^. Because dupα7 shares high sequence homology with α7, previous studies using non-specific primers may have overestimated α7 expression^11,20,67–76^. Our study provides the first clear evidence of dupα7 expression in these cell lines, establishing them as valuable models for studying the unique contributions of dupα7 to human T cell biology.

In Jurkat cells, 48 h of mitogenic stimulation induced a reciprocal shift in α7 and dupα7 expression, with dupα7 upregulated and α7 downregulated. Since dupα7 functions as a dominant-negative regulator of α7^50^, this shift may suppress α7-mediated signaling during prolonged T cell activation. Similar changes in the α7/dupα7 ratio have been reported in sepsis and inflammatory bowel disease^77,78^, supporting the relevance of our findings to human immune responses. We also observed downregulation of α9 and α10 expression following mitogenic stimulation, consistent with prior studies showing that TCR activation suppresses α9-containing nAChRs^29,79^. Since α9-containing receptors have high affinity for ACh^40,80^, their downregulation during activation may serve to fine-tune T cell cholinergic signaling.

Our functional assays highlight the complexity of cholinergic regulation on cytokine production. Whereas previous reports showed that pCF3-diEPP suppresses cytokine release in human whole blood cells^81^, it had no significant effect on IL-2 or TNF-α secretion in Jurkat cells, suggesting that nAChR signaling is cell type-dependent. Nicotine modestly inhibited cytokine production and significantly reduced %IL-2+ cells, consistent with its well-established immunosuppressive effects. Unexpectedly, antagonists of α7 (ArIB) and α9α10 (RgIA4) did not reverse the effects of nicotine but instead enhanced them. This contrasts murine T cell experiments, where nicotine-mediated IL-2 inhibition was reversed by an α7- and α9α10-selective antagonist^28^, and with macrophage studies where ArIB reversed α7-mediated anti-inflammatory effects^42,81^.

Interestingly, ArIB alone significantly inhibited IL-2 and TNF-α, suggesting that endogenous α7 signaling is required for robust cytokine production. This aligns with prior work showing that exogenous ACh enhances IL-2 release in rat T cells, an effect blocked by nAChR antagonists^82^, and with reports that α7 ligands suppress TNF-α in microglia^83^.

Differences between T cells and macrophages may reflect distinct regulation of the α7/dupα7 ratio. In macrophages, proinflammatory stimuli increase α7 relative to dupα7^84,85^, favoring homomeric α7 receptors that attenuate inflammation. In contrast, Jurkat cells showed decreased α7/dupα7 after stimulation, which may favor heteromeric α7dupα7 receptors with altered or reduced signaling capacity.

Another explanation lies in the distinct physiology of immune versus neuronal nAChRs^57^. In neurons, ligand binding opens ion channels, but in immune cells, nAChRs primarily signal via metabotropic protein-protein interactions, independent of ion flux^22^. Classical antagonists of neuronal α7 channel activity may therefore act as unconventional agonists in immune cells. Supporting this, the non-selective nAChR antagonist mecamylamine suppressed mitogen-induced IL-2 release in MOLT-3 T cells^29^. Although originally interpreted as evidence of endogenous ACh signaling, this could reflect the activation of non-canonical signaling pathways. Furthermore, several studies report that α7-mediated effects in immune cells persist despite conventional antagonists^38^, but are abrogated by α7 knockdown^83^ or tyrosine kinase inhibitors^23,67^. This may explain why the effects of nicotine were enhanced, rather than reversed, by ArIB or RgIA4 in our study.

The regulation of IL-2 by nAChR signaling has important therapeutic implications. While IL-2 supports effector T cell expansion, its primarily role is maintaining regulatory T cells (Tregs)^86^. Because Tregs express high-affinity IL-2 receptors, they outcompete effector T cells when IL-2 levels are low ^86^. Thus, nicotine-induced reductions in IL-2 may suppress effector T cell responses while supporting Treg expansion, tipping the balance toward immune tolerance. This is consistent with prior studies showing that nicotine enhances Treg development and suppresses effector T cell activity^28,33,87^. By selectively supporting Tregs, nicotine and α7-selective ligands such as ArIB may protect against autoimmune and inflammatory diseases.

Overall, our results support a model in which nAChR subtypes are dynamically regulated during T cell activation, directly influencing cytokine production and immune responses. The observed increase in dupα7 and decrease in α7 may represent a human-specific mechanism for tuning cholinergic immunoregulation. This shift could enable T cell-derived ACh to act preferentially on macrophages or other immune cells, promoting systemic control of inflammation.

This study has several limitations. All experiments were performed using immortalized T cell lines, which do not fully represent primary human T cells. Future work should validate these findings in primary T cells and in vivo models. CRISPR/Cas9-mediated knockout of specific nAChR subunits will also be essential to confirm their roles in cytokine regulation. Finally, elucidating the intracellular signaling pathways downstream of α7- and dupα7-containing receptors will be key to understanding how these receptors shape immune function.

In conclusion, Jurkat cells provide a robust model for studying nAChR expression and function in human T cells. Our findings highlight the previously overlooked role of dupα7 and demonstrate that endogenous α7-mediated signaling is essential for cytokine production. The dynamic regulation of α7 and dupα7 during activation may be a key mechanism for balancing inflammation and immune tolerance, offering potential therapeutic targets for autoimmune and inflammatory diseases.

## 8 Author Contributions

BL designed and performed experiments, analyzed data, and drafted the manuscript. AS supervised the study, provided resources, and edited the manuscript. All authors approved the final manuscript.

## Acknowledgments

We thank Dr. Sujeenthar Tharmalingam for providing qPCR training. We also thank Dr. Parissenti for generously providing ELISA kits and Dr. Roger Papke at the University of Utah for synthesizing and providing the silent agonists and conopeptides used in this project. Thank you to Dr. Jenny Bruin for providing feedback on the manuscript.

